# RNA-guided transcriptional repression by TIGR-Tas systems

**DOI:** 10.64898/2026.09.09.750476

**Authors:** Peiyu Xu, Lawrence Y. Long, Annie Zhu, Stephanie Kim, Shai Zilberzwige-Tal, Natalia Quinones-Olvera, Lilia Evgeniou, Daniel Flam-Shepherd, Rhiannon Macrae, Guilhem Faure, Feng Zhang

## Abstract

Tandem interspaced guide RNA (TIGR)–TIGR-associated protein (Tas) systems are a widespread family of RNA-guided DNA-targeting proteins whose diversity has remained uncharacterized because their arrays, unlike CRISPR arrays, lack the sequence conservation required by existing annotation tools. We developed TIGRFinder, a motif-based pipeline that identified 6,685 TIGR arrays from genomic and metagenomic data. Phylogenetic and structural analysis of Tas proteins revealed clade-specific insertions in stem-loop binding Tas proteins that co-vary with features of their cognate tigRNAs. Cryo-electron microscopy structures of two stem-loop binding TasR ribonucleoprotein complexes demonstrate how these protein insertions directly accommodate extended tigRNA stems while maintaining DNA binding through catalytically inactive RuvC domains. We also show that these catalytically dead TasR proteins, along with the nuclease-lacking TasA, function as RNA-guided transcriptional repressors.These findings establish TIGR-Tas as a functionally diverse family of RNA-guided effectors, with comprehensive annotations available through TIGRSafari, which will be made publicly available upon publication, to support further exploration and engineering.

## Main

TIGR-Tas (Tandem Interspaced Guide RNA – TIGR-Associated) systems are a recently described family of RNA-guided DNA-targeting proteins encoded predominantly by bacteriophages, archaeal viruses, and parasitic bacteria of the Candidate Phyla Radiation (CPR)^1^, identified through metagenomic sequencing Like CRISPR-Cas systems, TIGR-Tas systems encode guide RNAs in genomic arrays adjacent to their effector proteins, but the architecture of these arrays is fundamentally different. CRISPR arrays consist of identical short direct repeats separated by variable spacers, a regular structure that makes them straightforward to detect computationally^2,3^. TIGR arrays, by contrast, encode dual-spacer guide RNAs (tigRNAs) in which each typically 36nt functional unit contains two distinct spacers flanked by box C and box D motifs: these spacers act in tandem, with spacer A pairing to one strand of a DNA target and spacer B to the other, a mechanism with no equivalent in CRISPR systems. Two distinct array architectures encode these dual-spacer guides. In dual-repeat arrays, edge and loop repeats alternate with pairs of spacers in a regular but sequence-variable pattern. In stem-loop arrays, palindromic sequences flanking each spacer pair form hairpin structures defining tigRNA units that individually resemble snoRNA^1^. Tas proteins share a conserved Nop RNA-binding domain that recognizes tigRNAs for RNA-guided DNA targeting^1^. Beyond this core domain, there is substantial variation in Tas proteins; the Nop domain can be fused to an HNH nuclease (TasH), a RuvC nuclease (TasR), or present alone (TasA).

The full diversity of TIGR arrays and their relationship to Tas protein variation remain largely unexplored, in part because TIGR arrays cannot be detected by existing CRISPR annotation tools due to their distinct and variable architectures. Here we developed TIGRFinder, a suite of motif-anchored methods tailored to each array architecture, and applied it to over 20,000 *tas* loci to systematically annotate TIGR arrays and explore their diversity. Comparing tigRNA and Tas protein variation across this dataset, we uncovered extensive covariation between array architecture and protein structure, particularly within stem-loop systems. We investigated this covariation structurally and functionally, focusing on stem-loop binding Tas clades where both the tigRNA and protein have diversified beyond the conserved dual-repeat blueprint. Cryo-EM structures of two stem-loop binding TasR ribonucleoprotein (RNP) complexes from *Patescibacteria* group bacteria and *Candidatus Staskawiczbacteria* group bacteria reveal lineage-specific protein insertions that directly accommodate extended tigRNA stems, while catalytic residues in their RuvC domains are consistently inactivated. We show that these catalytically dead TasR proteins, along with the nuclease-lacking TasA, function as RNA-guided transcriptional repressors, revealing a previously unreported function of TIGR-Tas systems.

## Results

### TIGRFinder enables systematic discovery of TIGR arrays and reveals the global landscape of TIGR-Tas diversity

We investigated the TIGR-Tas diversity across two parallel axes: the organization of TIGR arrays and the diversity of Tas proteins. To characterize tigRNA diversity, we developed TIGRFinder, a computational tool for the systematic identification and annotation of TIGR arrays surrounding *tas* loci (Fig. 1a). TIGRFinder detects candidate tigRNAs using conserved motifs including box C and box D nucleotides flanking spacers as anchors for a seed-and-extend strategy, then iteratively relaxes constraints to recover degenerate repeats at array boundaries (see Methods). Separate sub-pipelines handle dual-repeat and stem-loop architectures, the latter requiring additional identification of palindromic stem sequences flanking each spacer pair (Fig. 1b). All annotations are made available through TIGRSafari (which will be made publicly available upon publication), an interactive web resource that integrates array annotations (including each tigRNA component) with Tas protein phylogeny and structural predictions.

**Figure 1.**
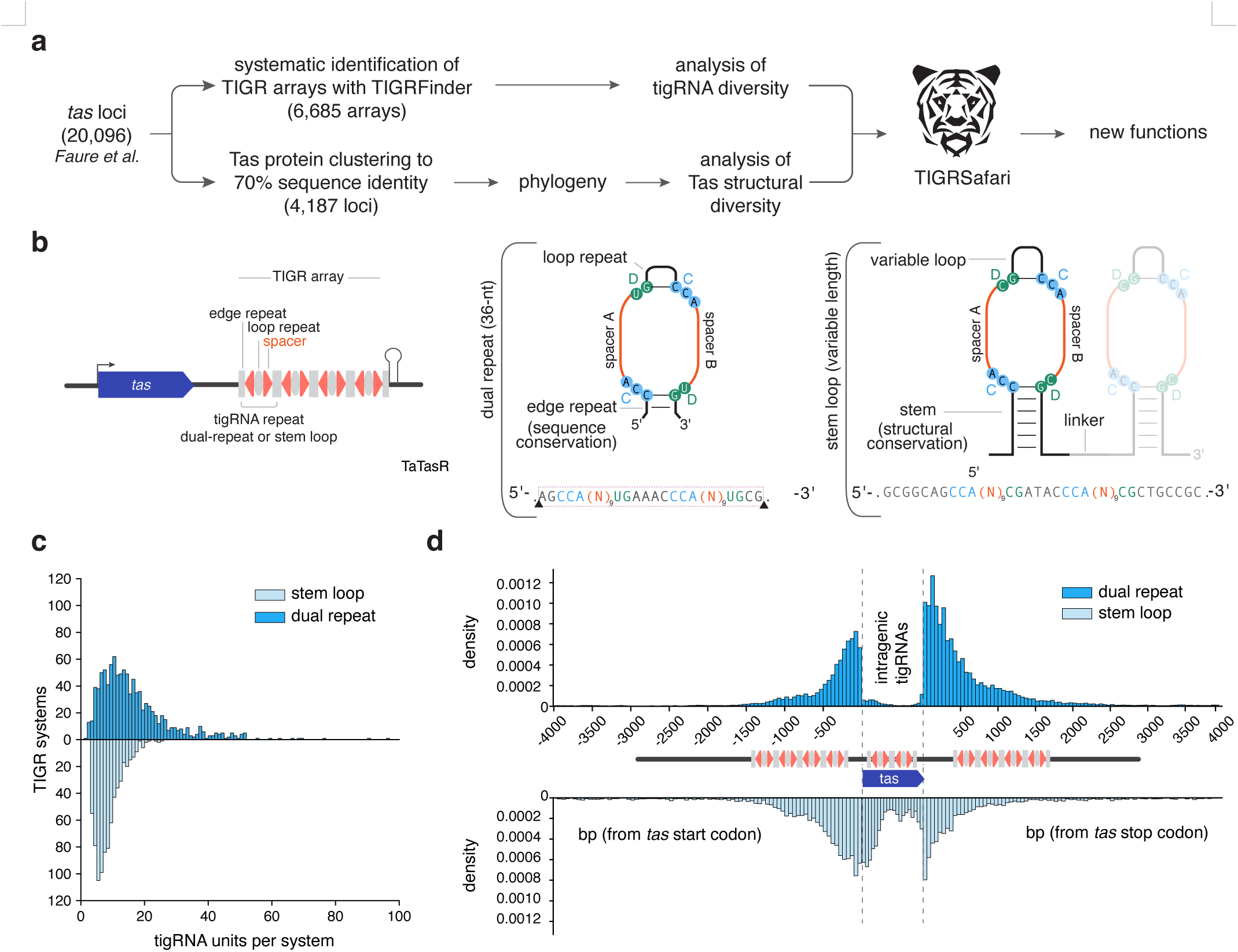
TIGRFinder catalogs diversity of TIGR systems. **(a)** Schematic of the analysis workflow. *Tas* loci were analyzed as follows: TIGR arrays were identified by TIGRFinder and Tas proteins were clustered for phylogenetic and structural analysis. All annotations are integrated into **TIGRSafari**. **(b)** Schematics of generalized TIGR-Tas array (left), dual-repeat tigRNA (middle), and stem-loop tigRNA (right). In the generalized schematic (left), repeats are shown in gray, spacers in red. Left and right facing triangles represent spacers A and B, respectively. Ovals show loop repeats, rectangles show edge repeats in dual-repeat systems and stems in stem-loop systems. For the tigRNA schematic (right), C box nucleotides are highlighted in cyan, D box nucleotides highlighted in green, spacers shown in red. **(c)** Distribution of TIGR units per TIGR-Tas system for **dual-repeat (DR)** and **stem-loop (SL)** architectures. The x-axis indicates the number of tigRNAs per system, and the y-axis indicates the number of TIGR-Tas systems. **(d)** Positional distribution of TIGR units relative to *tas*. Density plots show repeat locations upstream of the *tas* start codon and downstream of the *tas* stop codon for dual-repeat and stem-loop systems. TIGR units located within the *tas* coding region are shown on a normalized *tas* gene schematic below.

Applied to a set of 20,096 *tas* loci identified previously^1^, TIGRFinder uncovered 6,685 TIGR arrays comprising 65,635 tigRNAs from 4,126 dual repeat arrays and 17,756 tigRNAs from 2,559 stem-loop arrays. This represents a substantial expansion over the manually curated set reported previously^1^. Based on this extended set of systems, several trends were noticeable. First, dual-repeat loci typically encode longer arrays containing more tigRNA units per system than stem-loop systems (Fig. 1c), suggesting distinct constraints on array expansion and evolutionary adaptation between the two architectures. Second, TIGR arrays display flexible genomic location and can occur upstream, downstream, or flanking both sides of the *tas* gene (Fig. 1d). Additionally, stem-loop arrays (and less frequently dual repeat arrays) can be located within the *tas* coding sequence at the 5’ end, 3’ end, or internal regions, without disrupting the reading frame. In these cases, the intragenic tigRNAs typically correspond to disordered regions between secondary structure elements, preserving the integrity of the Nop domain fold. This integration of the guide RNA within the effector gene suggests tight co-evolution between the tigRNA and the Tas protein, and may link tigRNA maturation and Tas translation.

### Extensive structural diversification of stem-loop binding Tas proteins

We next investigated the diversity of Tas proteins themselves, focusing on the stem-loop clade where structural variation is most pronounced. We clustered non-redundant Tas sequences at 70% identity, a higher threshold than the 50% used previously^1^, to better resolve clade-level structural variation. We inferred a maximum-likelihood phylogeny using structurally aligned Nop domain positions (Fig. 2a, see Methods).

**Figure 2.**
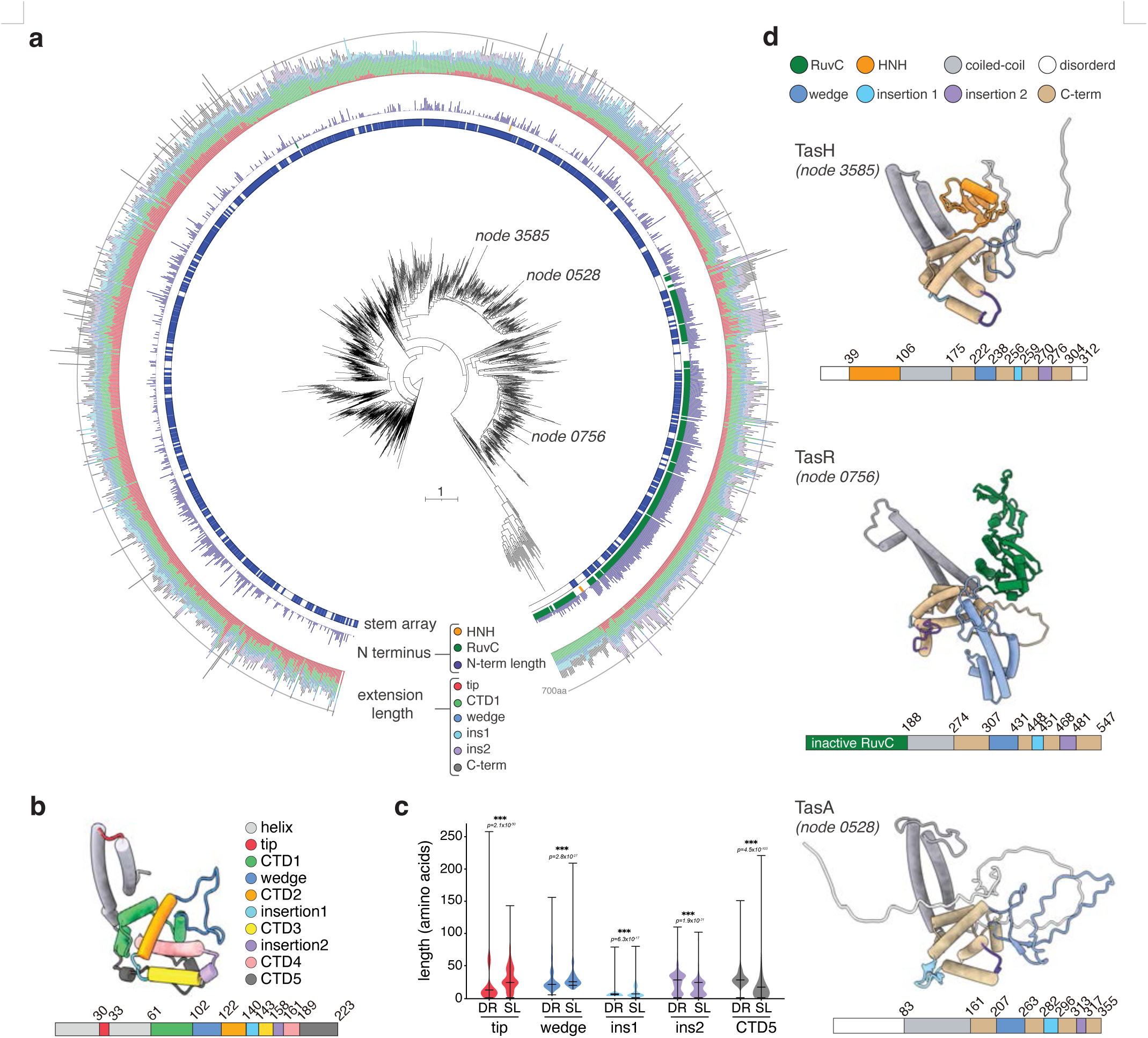
Diversity of Tas proteins. **(a)** Phylogenetic tree of stem-loop binding Tas protein representatives (clustered at 70% of sequence identity). Inner to outer rings show: stem-loop array detection by TIGRFinder, presence of nuclease (RuvC in green, HNH in orange), N-terminus Tas length, NOP domain extensions colored by their internal location shown in panel b (red for tip insertion, gray for coiled-coil, green for CTD1, dark-blue for wedge insertion, orange for CTD2, cyan for insertion1, yellow for CTD3, purple for insertion2, pink for CTD4, dark-gray for CTD5). Nodes of interest (see Fig. 2d) are labelled in the tree. Node IDs are mapped to full contig IDs in Supplementary Table. **(b)** Structural representation of the NOP domain colored by components (grey for coiled-coil domain, red for tip, green for CTD1, dark-blue for wedge insertion, orange for CTD2, cyan for insertion1, yellow for CTD3, purple for insertion2, pink for CTD4, dark-gray for CTD5). **(c)** Length distribution of Tas protein components, comparing dual-repeat binding Tas (DR) clade against stem-loop binding Tas (SL). Same color as in panel b. **(d)** AlphaFold models Tas proteins with varying N-terminal domains (node 3585 is TasH, node 0756 is TasR, node 0528 is TasA) indicated in the tree. N-terminal nuclease is shown in orange (HNH) or green (RuvC), coiled-coil is shown in grey, major insertions in dark-blue (wedge insertion), cyan (insertion1), and purple (insertion2), remaining components of NOP are in beige, disordered region outside NOP domain are shown in white.

The most variable feature across stem-loop binding Tas clades is the N-terminal region upstream of the Nop core. A large monophyletic clade composed predominantly of CPR bacteria (Extended Data Fig. 1a and Supplementary Table 1), the only known non-viral TIGR hosts with arrays, encodes TasR proteins with substantially expanded RuvC domains bearing extensive insertions (Fig. 2a). HNH-type nuclease domains appear at two phylogenetically isolated positions in the tree, while many remaining clades carry extended disordered N-terminal regions of varying length, as exemplified by TasA (Fig. 2a). One long phylogenetic branch lacked detectable TIGR arrays; manual inspection of the corresponding loci revealed compact TasR proteins lacking the coiled-coil dimerization domain, encoded within densely packed viral genomes with no apparent intergenic space to harbor an array (Extended Data Fig. 1a,b).

To systematically compare structural variation between dual-repeat and stem-loop systems, we decomposed the NOP core domain into defined segments: the coiled-coil helix tip, wedge, insertion 1 (between CTD2 and CTD3), insertion 2 (between CTD4 and CTD5), and the CTD5 region (Fig. 2b). We measured their length distributions across dual-repeat versus stem-loop associated Tas proteins (Fig. 2c). Dual-repeat binding Tas proteins maintain a compact NOP core with minimal variation across all segments (Extended Data Fig. 1a). Stem-loop-binding Tas proteins have significantly longer coiled-coil tips and insertion 1 regions, whereas insertion 2 is significantly longer in dual-repeat-binding Tas proteins (Fig. 2c). The precise pattern of expansion distributed non-uniformly across the phylogeny: specific clades display coordinated expansion at particular positions, suggesting lineage-specific structural adaptations (Extended Data Fig. 1a).

A critical exception to this trend is the CTD5 region, which is significantly shorter in stem-loop binding Tas compared to dual-repeat binding Tas (Fig. 2c). In dual-repeat systems, CTD5 harbors an extended loop containing conserved tyrosine and histidine residues that constitute a metal-ion-independent dyad catalytic site responsible for tigRNA processing^4^(Extended Data Fig. 1a). The consistent truncation of this region across stem-loop binding Tas clades indicates that these systems employ a distinct, as yet uncharacterized mechanism for tigRNA maturation, a hypothesis further supported by the identification of a conserved CUU processing motif at the stem base described below (Extended Data Fig. 1a).

Insertion 1, located between CTD2 and CTD3, binds within the interior of the tigRNA spacer loop, where it is thought to physically separate the two spacers and promote their accessibility for target recognition. Although frequently disordered, insertion 1 can adopt defined secondary structures in certain clades (Extended Data Fig. 2a and b), suggesting its conformation may be tuned to properties of the cognate tigRNA. Insertion 2, positioned between CTD4 and CTD5, is typically shorter and structurally heterogeneous, adopting beta-sheet, short helical, or disordered conformations depending on the clade (Extended Data Fig. 2b). This region has been hypothesized to function as a molecular ruler governing target interaction geometry^4^. Together, the diversity of these insertions reflects extensive structural elaboration in stem-loop binding Tas proteins, likely reflecting co-adaptation to their cognate tigRNAs or the acquisition of additional functions.

### Co-evolution of tigRNA architecture and Tas protein structure in stem-loop systems

The structural diversity of stem-loop binding Tas proteins is paralleled by remarkable variation in their cognate tigRNAs. Unlike dual-repeat tigRNAs, which adopt a conserved 36-nucleotide (nt) fold, stem-loop tigRNAs vary extensively in stem length, loop size, and the length of the linker between adjacent units (Fig. 1b; Extended Data Fig. 2a,e,f,g). Sequence conservation within the stem is largely restricted to a few nucleotides proximal to the box C and box D motifs, while the remainder of the stem diverges substantially even among tigRNAs within the same array (Extended Data Fig. 2c). Mapping these RNA structural features onto the Tas phylogeny reveals that loop length, linker length, and stem sequence conservation vary in a clade-specific manner (Extended Data Fig. 2a), consistent with co-evolution between each Tas protein and its cognate guide RNA architecture. While the loop between the two spacers are mostly conserved to be 9 nt long similar to dual-repeat (Extended Data Fig. 2e, Extended Data Fig. 1a), in some cases for stem-loop clades, the loop between the two spacers folds into a secondary hairpin structure (Extended Data Fig. 2d), adding a further layer of structural complexity. For instance, the CPR phyla TasR clade harbors a large conserved insertion along with the tigRNA having an extended hairpin located in the loop region (Extended Data Fig. 2a). The TasA proteins carry an extended CTD5 domain, which might indicate an alternate processing mechanism of tigRNA maturation in this specific clade of TIGRs. Overall, stem-loop TIGR systems exhibit conserved combinations of tigRNA traits that track with conserved structural modules in Tas proteins, indicating coordinated co-evolution of the guide RNAs and their associated Tas effectors.

This coordinated diversity in both the protein and RNA components of stem-loop TIGR systems motivated us to determine experimental structures of several stem-loop binding Tas RNP complexes. We focused on two systems representing distinct phylogenetic clades with contrasting structural predictions: *Patescibacteria* group bacterium TasR (PateTasR), which has a compact RuvC domain and relatively short tigRNA stems, and *Candidatus Staskawiczbacteria* bacterium TasR (CsTasR), a CPR-derived protein with a substantially expanded RuvC domain predicted to accommodate extended tigRNA stems.

### Cryo-EM structure of PateTasR reveals a catalytically inactive RuvC domain co-adapted to extended tigRNA stems

We purified PateTasR (329aa) co-expressed with its native TIGR array from *E. coli* and determined its cryo-EM structure (Fig. 3a, Extended Data Fig. 3) . The complex adopts a C2- symmetric dimer mediated by a coiled-coil scaffold, with an X-shaped architecture in top view and a 2:1 stoichiometry of TasR to tigRNA, similar to the TaTasR structure (PDB: 9MTY)^1^. A characteristic kink in the upper coiled-coil helix positions the two RuvC domains symmetrically on opposing faces of the dimer (Fig. 3a). The coiled-coil stabilizes RuvC helices α1 and α2, stabilizing the nuclease-like fold through an extensive interdomain interface. Specifically, D114 forms a salt bridge with R27 of the RuvC domain, while R224 engages in a stacking interaction with Y108 (Fig. 3b); on the opposing interface, E161 from the coiled-coil coordinates with R58 and R32 of the RuvC domain (Fig. 3c). Together, these interactions rigidly anchor the RuvC domain to the coiled-coil scaffold, resulting in a more compact overall architecture than the dual- repeat TaTasR.

**Figure 3.**
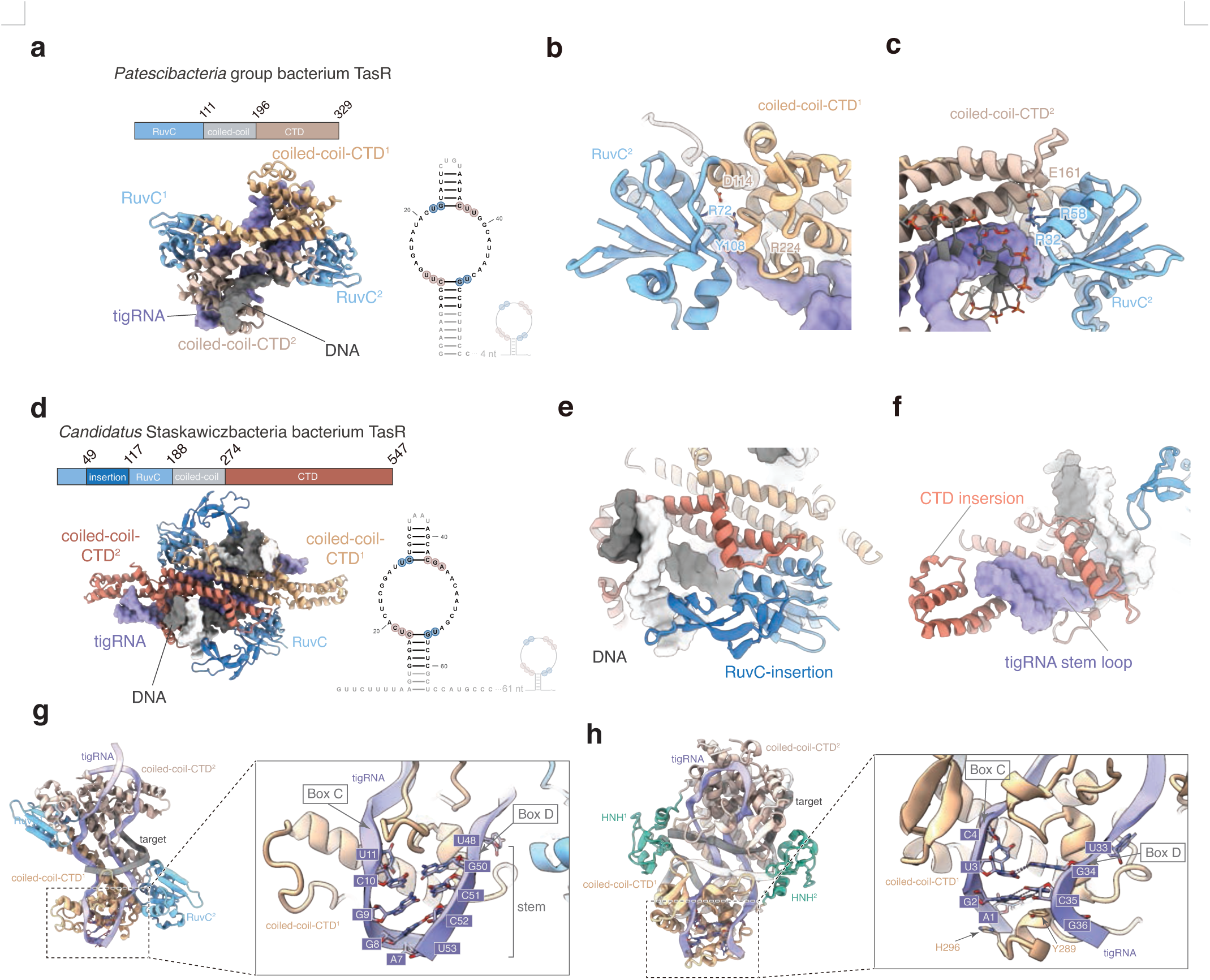
Cryo-EM structures of *Patescibacteria* group bacterium TasR and *Candidatus Staskawiczbacteria bacterium* TasR. **(a)** Domain architecture (top) and cryo-EM structure (bottom) of the Patescibacteria group bacterium TasR (PateTasR) complex (2.8 Å resolution) shown with the secondary structure prediction of a tigRNA (box C colored in tan and box D colored in blue). Color coding: RuvC domain, blue; coiled-coil scaffold, gray; C-terminal domain (CTD), tan; tigRNA, purple; target substrate, black. **(b)** Detailed view of the interaction interface between the coiled-coil-CTD1 of one protomer and the RuvC domain of the opposing protomer (RuvC2). Key interacting residues are shown as sticks and labeled. **(c)** Close-up view of the structural interactions between coiled-coil-CTD2 and RuvC2. Proteins are represented as cartoon models, with specific residues highlighted as sticks. **(d)** Domain architecture (top) and cryo-EM structure (bottom) of the *Candidatus Staskawiczbacteria bacterium* TasR (CsTasR) complex. Color coding: RuvC domain, light blue; RuvC insertion (80 amino acids), dark blue; coiled-coil scaffold, gray; CTD, tan; tigRNA, purple; target substrate strand 1, black; target substrate strand 2, white **(e)** Structural detail showing the RuvC insertion positioned proximal to the DNA substrate. The DNA and protein residues are shown as sticks. **(f)** Close-up view highlighting the CTD insertion as it wraps around the stem region of the tigRNA. **(g)** The structure of the stem region in the PateTasR complex. Left: overall structure in top view. Right: close-up view of the Box C/D and stem regions of the tigRNA; the protein is shown as a cartoon, while Box C/D and base-pairing bases are represented as labeled sticks. **(h)** The structure of the tigRNA in the dual-repeat binding TasH (PDB: 9W04) ^4^. Left: overall structure in top view; the protein is shown as a cartoon, with Box C/D and base-pairing bases represented as labeled sticks. Right: close-up view of the Box C/D region.

The PateTasR tigRNA adopts a figure-eight conformation separated by the Tas wedge loop, topologically similar to the 36-nt tigRNA of TaTasR, despite being longer (∼54 nt) (Extended Data Fig. 4a and b). The box C/D motif of PateTasR consists of two symmetric CUU/UG sequences. A base flip occurs between the two uracils of box C, which are sandwiched between the wedge loop and the terminal helix of the C-terminal domain (Extended Data Fig. 4c). The cytosine of box C forms a base pair with the guanine of box D, while the first U engages N315 via hydrogen bonding and the A makes a stacking interaction with Y304 (Extended Data Fig. 4d). Notably, the U of box D interacts with R225 in the CTD and S75 in the RuvC domain—a different interaction geometry from dual-repeat TaTasR, where R198 at the C-terminus of the RuvC domain stabilizes a uracil at box D (Extended Data Fig. 4c and d). This structural divergence in box D recognition likely dictates the protein’s specificity for cytosine versus uracil at this position.

Examination of the RuvC active site reveals that PateTasR is catalytically inactive: the canonical catalytic residues are absent, and R91 sterically occupies the catalytic pocket, neutralizing any residual charge and precluding substrate cleavage. Consistent with this, the C-terminal loop extending beyond the final CTD helix is remarkably short in PateTasR, comprising only 8 amino acids compared to the 27-amino-acid extension in SpTasH that harbors the Y/H catalytic dyad for tigRNA processing (Fig. 3g and h)^4^. The absence of this bulky loop extension creates sufficient space to accommodate the extended distal stem of the stem-loop tigRNA, revealing a structural solution to the geometric demands of longer guide RNAs. Despite lacking cleavage activity, we observed unassigned density consistent with a single-stranded DNA target in the RNP complex, spanning 13 nucleotides modeled based on the spacer sequence, demonstrating that PateTasR retains DNA-binding capacity through surrounding electrostatic and hydrogen-bonding interactions (C208, Q131, K10, T254, G252, R250) that collectively stabilize the substrate backbone at both ends and in the middle (Fig. 3 and Extended Data Fig. 4e-g). The DNA substrate is away from the inactive catalytic pocket (Extended Data Fig. 4h-i).

### The CsTasR structure reveals co-adaptation of expanded protein insertions with extended tigRNA stems

To capture the structural diversity at the other end of the stem-loop spectrum, we expressed and purified CsTasR (547aa), the CPR-derived TasR with a predicted extended RuvC insertion, and determined its cryo-EM structure in the presence of a 20-bp dsDNA target (Fig. 3d). Not only is CsTasR larger than PateTasR, but the predicted cognate tigRNA possesses extended terminal stems and two 10-nt spacers (Fig. 3d). Single-particle analysis resolved the structure of a DNA-bound RNP complex at 3.3 Å resolution (Extended Data Fig. 5a-f).

CsTasR adopts a C2-symmetric dimer mediated by a coiled-coil scaffold with overall resemblance to PateTasR, exhibiting a 2:1:1 stoichiometry of TasR:tigRNA:DNA (Fig. 3d). However, the RuvC domain of CsTasR contains an ∼80-residue insertion between strands β3 and β4, unique among characterized Tas proteins and absent in dual-repeat TaTasR, Cas12, TnpB, and Fanzor^1,5–8^. This insertion folds into a small subdomain comprising a short α-helix, a curved β-sheet, and loop-rich elements positioned proximal to the target DNA. (Fig. 3e).

Additionally, the C-terminal domain of CsTasR features a substantial insertion that wraps around the stem region of the tigRNA, yielding continuous EM density for this stem segment and suggesting steric protection by the protein (Fig. 3f). This structural arrangement implies a direct co-adaptation between the CTD insertion and the tigRNA stem linker extension, an RNA-protein interface that is absent in systems with shorter stems.

As with PateTasR, the RuvC domain of CsTasR is catalytically inactive: the first two canonical catalytic positions are filled by R8 and V126 (Extended Data Fig. 5g). In the fully engaged state, each spacer pairs with only 9 nucleotides of the target, demonstrating that CsTasR, like its relatives, recognizes an 18-bp target overall. Together, the two structures demonstrate a coherent pattern: stem-loop binding Tas proteins have evolved RuvC domains that retain DNA-binding scaffolding while losing nuclease activity, and have accumulated lineage-specific insertions in the NOP regions that co-evolve structurally with the extended stems of their cognate tigRNAs.

### Stem-loop binding Tas proteins lack the Y/H processing loop found in dual-repeat systems

Dual-repeat binding TasH processes precursor tigRNAs into mature 36-nt guides through a metal-ion-independent mechanism mediated by conserved tyrosine and histidine (Y/H) residues located on an extended loop at the C-terminus of the NOP domain^4^. Sequence alignments across Tas proteins show that this Y/H dyad is conserved in dual-repeat systems but absent across all stem-loop clades, including the rare stem-loop binding TasH proteins found at only two isolated positions in the phylogeny (Fig. 2a; Extended Data Fig. 1a). Consistent with this, both PateTasR and CsTasR lack this 27-residue C-terminal catalytic loop (Fig. 3g, h). In PateTasR, the region is reduced to just 8 amino acids, leaving space for the extended distal stem of stem-loop tigRNAs but without the catalytic machinery for tigRNA processing. The absence of Y/H across tem-loop systems regardless of N-terminal domain architecture, combined with the extended distal stems of stem-loop tigRNAs, suggest that stem-loop TIGR systems employ a distinct tigRNA maturation mechanism.

### Stem-loop tigRNAs are expressed from flexible genomic positions and processed at a conserved CUU motif

To characterize tigRNA expression and processing in stem-loop binding Tas systems experimentally, we co-expressed CsTasR and *Bifidobacterium longum* TasA (BlTasA) with their respective native loci in *E. coli* and then purified the resulting RNP complexes, extracted RNA, and performed small RNA sequencing (smRNA-seq). Both loci produced multiple processed tigRNA units (Fig. 4a and b).

**Figure 4.**
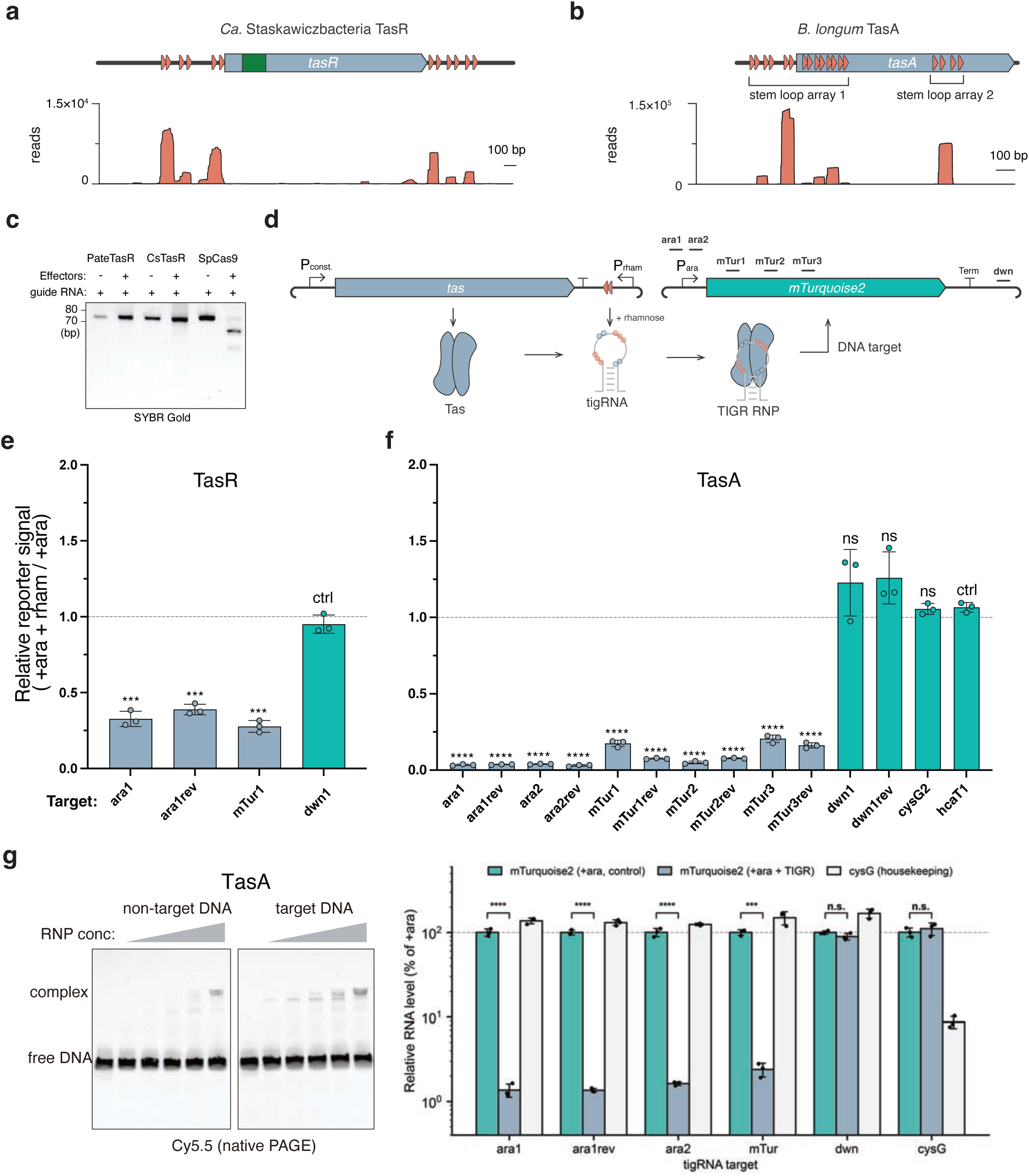
RNA sequencing and DNA targeting of stem-loop binding TasR and TasA. **(a)** Locus architecture (top); small RNA sequencing profile (bottom) of the *Candidatus Staskawiczbacteria bacterium* TIGR. **(b)** Locus architecture (top); small RNA sequencing profile (bottom) of the *Bifidobacterium longum* TIGR. **(c)** In vitro nuclease assay of PateTasR and CsTasR complexes with target DNA. The 2% agarose gel shows DNA migration following proteinase treatment and column purification. **(d)** Schematic of the in vivo reporter assay for TIGR-mediated transcriptional interference in E. coli. Components include: arabinose-inducible tas genes, T7-driven tigRNAs, and a rhamnose- inducible mTurquoise reporter. TS1–TS3: target sites 1–3. **(e, f)** mTurquoise fluorescence quantification for *Candidatus Staskawiczbacteria bacterium* TasR (e) and *Minisyncoccota bacterium* TasA (f). ’Control’: co-expression of the Tas protein with its native, non-targeting tigRNA. ’TS1/2/3’: co-expression with reprogrammed tigRNAs targeting specific sites within the mTurquoise gene. **(g)** Electrophoretic mobility shift assay (EMSA) of the purified TasA RNP with target DNA and a non-target DNA control. The DNA substrates are 5’-labeled with Cy5.

CsTasR tigRNAs were found both upstream and downstream of the protein-coding sequence, while BlTasA tigRNAs were found not only upstream but also embedded within the coding sequence itself (Fig. 4a and b). AlphaFold predictions indicate that the intragenic tigRNA segment in BlTasA maps to the wedge loop domain of the protein, a domain that directly contacts the tigRNA in the mature complex. This spatial correspondence raises the possibility that the wedge loop ancestrally co-evolved with an embedded guide RNA, potentially coupling tigRNA maturation with Tas translation (Extended Data Fig. 5h). Secondary structure analyses confirmed the tigRNA architectures inferred computationally: CsTasR tigRNAs possess a 5-bp stem between spacers, consistent with the CTD insertion that contacts this stem in the cryo-EM structure (Fig. 3d). Processed tigRNA boundaries frequently coincide with a conserved CUU trinucleotide motif (Extended Data Fig. 6a-c). This motif is highly conserved across diverse stem-loop tigRNAs (Extended Data Fig. 6d) and consistently maps to the base of the stem structure (Extended Data Fig. 6e). Notably, tigRNAs co-expressed with extended flanking sequences, including 2,215 nt of downstream flanking sequence adjacent to the TIGR locus, exhibited sharper, more defined transcript boundaries and greater RNA uniformity on denaturing gels—suggesting that upstream or downstream sequence context facilitates processing (Extended Data Fig. 6c). Together, these data implicate that while the conserved CUU motif defines the specific site for tigRNA processing, elements within the extended flanking sequences function to facilitate and enhance this processing. This stem-loop biogenesis model is distinct from the dual-repeat mechanism, in which the Tas Y/H catalytic dyad cleaves within the conserved edge repeat to release mature 36-nt tigRNAs.

### Catalytically inactive TasR and TasA proteins function as RNA-guided transcriptional repressors

Both PateTasR and CsTasR harbor RuvC domains that appear catalytically inactive based on structural analysis, with substitutions at canonical positions. To confirm this experimentally, and test if these proteins, as well as a distinct TasA from the *Minisyncoccota bacterium* (MbTasA), nonetheless bind DNA, we performed in vitro nuclease assays with purified RNPs and Cy5- labeled dsDNA substrates. No detectable DNA cleavage was observed for PateTasR,or CsTasR, consistent with the inactivated RuvC domains of the TasR proteins and the absence of any nuclease domain in TasA (Fig. 4c).

To test DNA binding in a cellular context, we designed an *in vivo* reporter assay in which the Tas protein and its cognate tigRNA were co-expressed with a reporter plasmid harboring an *mTurquoise* gene targeted by the tigRNA (Fig. 4d). Co-induction of the TIGR system and the reporter gene led to strong repression of *mTurquoise* expression for both CsTasR and MbTasA (Fig. 4e and f). Repression was tigRNA sequence-dependent: co-expression with non-targeting native tigRNAs produced no significant reduction in fluorescence. This is consistent with the RNP complex binding the target site and sterically blocking transcription. Withdrawal of rhamnose after tigRNA induction restored mTurquoise2 fluorescence for both CsTasR and MbTasA, whereas cultures maintained on rhamnose remained repressed (Extended Data Fig. 7c-d), indicating that Tas-mediated transcriptional repression is reversible rather than a consequence of irreversible target loss. RT-qPCR confirmed that repression is RNA-guided and target-specific, with tigRNAs directed against the mTurquoise2 reporter reducing its transcript while leaving host housekeeping genes unchanged. Redirecting the tigRNA to an endogenous E. coli gene (cysG) likewise repressed that transcript, showing that TIGR-Tas can silence native chromosomal genes (Fig. 4h).

To confirm DNA-binding of MbTasA *in vitro*, we performed electrophoretic mobility shift assays (EMSA) with purified RNP and Cy5-labeled dsDNA substrates. The TasA RNP exhibited high-affinity binding to target DNA relative to a non-target control, producing a complete mobility shift at low RNP concentrations (Fig. 4g). Together, these results demonstrate that catalytically inactive TasR proteins and TasA function as RNA-guided DNA-binding modules capable of sequence-specific transcriptional repression.

## Discussion

A central theme emerging from this work is the coordinated co-evolution of Tas protein architecture and tigRNA structure in stem-loop systems. While dual-repeat TIGR systems maintain a conserved NOP domain, and uniform 36-nt guide RNA, stem loop clades have diversified extensively: conserved combinations of protein insertions and tigRNA structural features across multiple lineages indicate that the two components have diversified in a coordinated manner. The CPR clade illustrates the diversity of Tas insertion and tigRNA features, harboring a conserved RuvC domain alongside some of the largest insertions observed both in the RuvC and NOP domains. Structural analysis reveals that these insertions serve distinct roles: although catalytically dead, the RuvC domain retains strong DNA interaction within the R-loop, with certain insertions extending its binding to the flanking dsDNA regions; NOP insertions in turn, contact extended tigRNA features including the stem and loop hairpins. The consistent absence of the Tas C-terminal processing loop across all stem loop clades reflects a fundamental geometric constraint: dual-repeat tigRNAs have a fixed 36-nt length that positions the cleavage site within reach of the Y/H catalytic dyad, but stem-loop tigRNAs form stems of variable length, displacing the maturation site far from the protein surface. This variability precludes the fixed protein-based cleavage mechanism used in dual-repeat systems. Instead, a conserved CUU trinucleotide motif at the stem base marks the processing boundary, though whether cleavage is carried out by the Tas protein through an as yet unidentified mechanism, by host ribonucleases, or by auxiliary factors remains to be determined. Across the broader dataset, genomic sequence alone cannot establish whether tas and tigRNAs are co-transcribed or expressed from separate promoters. Intragenic tigRNAs could arise either through processing of a longer tas-containing transcript or through independent transcription from one or more promoters embedded within tas, and these mechanisms may differ among TIGR-Tas lineages. Our small RNA sequencing shows that TIGR-Tas arrays are processed into discrete tigRNAs even in the heterologous host *E. coli*, and because our ligation-free workflow preserves native 5′ ends, we could define mature tigRNA boundaries with confidence. That maturation occurs at all in E. coli suggests that processing relies on a conserved or host-supplied RNase rather than a dedicated system-encoded nuclease. However, our data do not report tigRNA 5′-phosphorylation state and thus cannot distinguish processing of a longer precursor from independent, promoter-driven transcription, particularly for internal segments such as stem-loop array 2, a question that will require end-chemistry–selective assays or transcription-start-site mapping to resolve.

Our biochemical results demonstrate that catalytically inactive Tas proteins function as RNA-guided transcriptional repressors. Both TasR variants we characterized carry substitutions in the RuvC catalytic residues, and TasA lacks a nuclease domain entirely; yet all bind target DNA with high affinity and silence gene expression in vivo. Consistent with an RNA-guided target-search mechanism, the RNP engages non-target DNA only weakly and at high concentrations, but binds target DNA at substantially lower concentrations, indicating a clear preference for the target sequence. On target DNA the complex resolves into two shifted species, in agreement with our cryo-EM structure showing that the effector assembles as a dimer, such that sequential occupancy of a single target by both protomers gives rise to the higher-order complex. The inactivated nuclease scaffold, rather than being a non-functional relic, has been repurposed to provide stable, guide RNA-directed DNA binding sufficient to block transcription. This natural transition from nuclease to repressor parallels the engineering of catalytically dead Cas9 as a programmable gene silencer^9^ ^10^, as well as the naturally inactive Cas12k, which retains its RuvC fold while acquiring a new function in guiding Tn7-like transposon insertion ^11^. TIGR systems thus represent a novel class of naturally occurring, nuclease-deficient RNA-guided DNA binding. The prevalence of TIGR systems in phages and CPR bacteria suggests that this repressor activity may serve in host gene silencing during infection, suppression of competing mobile elements, or regulation of the viral lifecycle. Determining which of these roles predominates will require functional studies in native host contexts.

Several fundamental questions remain. How TIGR arrays acquire new spacers is unknown: no adaptation machinery has been identified, and the architecture of tigRNAs imposes strong constraints on how new spacers working in tandem can be incorporated into a tigRNA unit while satisfying both structural and sequence requirements. The natural targets of most spacers cannot yet be determined, as the majority of TIGR systems reside on metagenomic contigs lacking broader genomic context, and the short spacer length generates a large number of false positive target matches. TIGRFinder successfully annotated arrays and components across most Tas loci, but at least one divergent branch lacked detectable arrays entirely, encoding compact TasR proteins within densely packed viral genomes with no intergenic space for an array. Whether these represent arrayless systems with an alternative guide RNA source, or simply escape current detection methods, is an open question. All annotations generated in this study are available through TIGRSafari (which will be made publicly available upon publication), providing a foundation for continued exploration and functional characterization of these diverse RNA-guided systems.

## Methods

### Cloning

Plasmids used in this study were cloned by Gibson assembly with NEBuilder HiFi DNA Assembly Master Mix (New England Biolabs, E2621L) and KLD Enzyme Mix (New England Biolabs, M0554S). The Stbl3 *E. coli* strain (Thermo Fisher, C737303) was used for DNA cloning. A N-terminal His-maltose-binding protein (MBP) tag was inserted after the start of Tas. Construct sequences were confirmed by whole-plasmid sequencing following Tn5 tagmentation after mini-prep of plasmids^12^ with QIAprep reagents (QIAGEN, 27106).

### TIGR RNP expression and purification

TIGR orthologs were expressed in *E. coli* T7 express cells (New England Biolabs, C2566H) and MBP-affinity purified. The expression vector was transformed into a T7 Express Competent *E. coli*. Colonies were transferred into a 10-mL starter culture of LB media for 17 hours, which was used to inoculate 1 L of TB media supplemented with 50 µg/ml spectinomycin and 50 µg/ml kanamycin for growth at 37°C and shaking at 180 rpm until an OD600 of 0.8 was reached. Protein expression was induced in the presence of 0.5 mM IPTG for 16 hours at 16°C. The cells were harvested by centrifugation for 10 minutes at 4°C at 4000 rpm (Beckman Coulter Avanti J-E, rotor JLA8.100). The pellet was kept frozen at −80°C for further use.

All purification steps were performed at 4°C. The *E. coli* cell pellet was resuspended in a buffer containing 20 mM HEPES pH 7.5, 150 mM KCl, and 4.5 mM TCEP supplemented with EDTA-Free Protease Inhibitor Cocktail (MedChem Express HY-K0010). The cell suspension was then subjected to a microfluidizer for lysis. The cleared lysate was applied to 3 mL of packed Amylose Resin (NEB) for 3 hours, followed by washing with 100 mL of buffer containing 20 mM HEPES pH 7.5, 150 mM KCl, and 4.5 mM TCEP. The MBP-tagged RNPs were then eluted with the same buffer supplemented with 10 mM Maltose (Sigma-Aldrich, M9171) and concentrated using an Amicon Ultra-15 Centrifugal Filter Unit (50KDa NMWL, Millipore UFC905024). The resulting elution was tested for the presence of the protein by NuPAGE (Invitrogen) and eStain L1 Protein Staining System (GenScript). For testing the presence of RNA, 1 µl Proteinase K (NEB) was mixed with 15 µl eluted RNP for 15 minutes and evaluated by 15% TBE-Urea polyacrylamide gels (Thermo Fisher Scientific).

### Small RNA sequencing

Purified RNPs (100 µL) were mixed with 600 µL of TRI Reagent (Zymo Research) and incubated for 5 min at room temperature. Chloroform (120 µL; Sigma-Aldrich) was added, and the sample was mixed by gentle inversion, incubated for 3 min at room temperature, and centrifuged at 12,000 × g for 15 min at 4 °C. The aqueous phase was used as input for RNA extraction with a RNA Clean & Concentrator-5 kit (Zymo Research), including on-column DNase I treatment, according to the manufacturer’s instructions. RNA was purified with an RNA Clean & Concentrator-5 kit (Zymo Research). Sequencing libraries were prepared using a SMARTer smRNA-Seq Kit for Illumina (Takara Bio) according to the manufacturer’s instructions, without any prior enzymatic modification of RNA termini. In this workflow, RNAs are polyadenylated at their 3′ ends, reverse-transcribed with an oligo(dT)-based primer, and the 5′ adapter is incorporated by template switching. Because library construction is ligation-free and does not require a 5′-monophosphate, tigRNAs were captured irrespective of their 5′-end chemistry (5′- hydroxyl, 5′-monophosphate, or 5′-poly/triphosphate), preserving their native 5′ ends; capture instead requires a free 3′-hydroxyl for poly(A) tailing. Libraries were sequenced on an Illumina MiSeq (Read 1, 80 cycles; Index 1, 8 cycles; Index 2, 8 cycles). The first three nucleotides added by template switching and the 3′ poly(A) sequence/adapters were removed with Cutadapt ^13^, and trimmed reads were mapped to the loci of interest with Bowtie 2 ^14^.

### In vitro cleavage assays

Double-stranded DNA (dsDNA) substrates were produced by PCR amplification of plasmids containing the target sites. Target cleavage assays were performed in a 10 µl reaction mixture containing 20 ng of substrate, 1 µM of TasR RNP or SpCas9 as a control in a final 1x reaction buffer of 20 mM HEPES pH 7.5, 150 mM KCl, and 5 mM MgCl2. Assays were allowed to proceed at 37°C for 2 hours. Reactions were then treated with RNase A (Qiagen) and Proteinase K (NEB) and purified using a PCR cleanup kit (Qiagen). Purified DNA substrates products were resolved by gel electrophoresis on E-gel 2% (dsDNA substrates), 15% TBE-Urea polyacrylamide gels (Thermo Fisher Scientific).

### TIGR gene repression assays

*Candidatus Staskawiczbacteria bacterium* TasR and *Minisyncoccota bacterium* TasA stem-loop TIGR were used for TIGR gene repression assays. To quantify the efficiency of RNA-guided transcriptional repression mediated by the TIGR-Tas system in vivo, a dual-inducible fluorescent reporter assay was constructed. The reporter gene mTurquoise2 (a cyan fluorescent protein variant) was cloned into a pCDF plasmid backbone under the control of an arabinose-inducible promoter. The Tas effector proteins (TasA or TasR) and their target-specific tigRNAs were carried on a separate effector plasmid derived from pEcCas (Addgene #73227): the effector protein was constitutively expressed from the native promoter (replacing Cas9), and the tigRNA was placed under the control of the rhamnose-inducible PrhaB promoter (replacing the sgRNA). The arabinose-inducible λ-Red recombination module was removed from this backbone to eliminate crosstalk with the arabinose-inducible reporter; the pEcCas backbone carries a non-temperature-sensitive replicon, allowing stable maintenance at 37 °C.

The reporter plasmid was first transformed into E. coli BW25113, a K-12 derivative carrying deletions of both the arabinose (ΔaraBAD) and rhamnose (ΔrhaBAD) catabolic operons; because this host cannot metabolize either inducer, arabinose and rhamnose accumulate to defined, non-depleting intracellular concentrations, providing tight and reproducible dose control. The resulting strain was rendered chemically competent using the Mix & Go! E. coli Transformation Kit (Zymo Research) according to the manufacturer’s instructions, and subsequently transformed with the effector plasmid. Co-transformants were cultivated in Luria-Bertani (LB) medium supplemented with the appropriate antibiotics (spectinomycin and kanamycin) and grown overnight at 37 °C with shaking at 220 rpm. The following day, 5 µL of the overnight culture was inoculated into 95 µL of fresh LB medium containing the corresponding antibiotics, arabinose (to induce mTurquoise2 expression), and rhamnose (to induce tigRNA expression). Cells were grown to mid-logarithmic phase at 37 °C. Culture fluorescence was measured using a BioTek microplate reader at an excitation wavelength of 434 nm and an emission wavelength of 477 nm. Data analysis was performed using GraphPad Prism 10. All experiments were performed with at least three independent biological replicates (triplicate cultures), and data are presented as the mean relative fluorescence units (RFU) with error bars representing the standard deviation (SD).

### Electrophoretic mobility shift assay (EMSA)

DNA binding by the MbTIGR–Tas ribonucleoprotein (RNP) complex was assessed using double-stranded DNA (dsDNA) probes. A target probe containing the tigRNA target sequence and a non-target probe were prepared by annealing equimolar complementary oligonucleotides (IDT), one strand of which carried a 5’ Cy5.5 fluorescent label; strands were heated to 95 °C for 3 min in a buffer containing 20 mM HEPES, 150 mM KCl, and 5 mM MgCl₂, and then cooled slowly to room temperature. The MbTIGR–Tas RNP was purified by size-exclusion chromatography (SEC), and the RNP peak fractions were pooled. Binding reactions (16 µl) contained 36 ng of Cy5.5-labeled dsDNA probe (1 µl of a 36 ng/µl stock) and a two-fold serial dilution series of SEC-purified RNP (15 µl), with the pooled peak fraction used undiluted as the highest concentration, in 20 mM HEPES, 150 mM KCl, 5 mM MgCl₂, and 15% glycerol. Reactions were incubated at 37 °C for 30 min. Binding reactions did not include a non-specific competitor; accordingly, the residual shift of non-target DNA observed at the highest RNP concentrations reflects general nucleic-acid affinity rather than guide-independent target recognition. Then 5 µl of TBE loading buffer was added, and 15 µl of each reaction was resolved on a 10% native TBE polyacrylamide gel run at 100 V for ∼120 min in 0.5× TBE at 4 °C. Gels were imaged in the Cy5.5 channel on a BioRad ChemiDoc. Free and RNP-bound DNA were detected as the “Free DNA” and “Shifted” (complex) bands, respectively.

### Cryo-EM data collection and processing

The prepared grids were transferred to the EF-Krios (Thermo Fisher Scientific) operating at 300 kV with a GatanK3 imaging system collecting at 105,000x nominal magnification. The calibrated pixel size of 0.4125 Å was used for processing. Movies were collected using Leginon 3.6 ^15^. Data were collected at a dose rate of 25.13 e-/Å2/s for PateTasR and 26.23 e-/Å2/s for CsTasR. Intermediate frames were recorded every 0.03 seconds for a total of 60 frames per micrograph. A total of 2,898 images for PateTasR and 6,284 images for CsTasR were collected at a nominal defocus range of 0.5 – 2.5 μm.

Image processing was performed on CryoSPARC v4.2.0^16^ and RELION 4.0^17^. Image stacks were subjected to beam-induced motion correction using MotionCor2.0^18^. Contrast transfer function (CTF) parameters for each non-dose-weighted micrograph were determined by CTFFIND4^19^. On-the-fly particle picking was done by Warp^20^.

For PateTasR, initial picking yielded 1,099,696 particles. Following 7 rounds of heterogeneous refinement and 1 round of 3D classification, we isolated 34,352 particles with well-defined structural features of a ternary complex. Subsequent non-uniform refinement^21^ generated a map with an indicated global resolution of 2.9 Å at a Fourier shell correlation (FSC) of 0.143. For CsTasR, initial picking yielded 1,455,905 particles. After 8 rounds of heterogeneous refinement, 125,754 particles showing well-defined structural features of the complex were retained. Non- uniform refinement^21^ of this subset produced a map with a global resolution of 3.3 Å (FSC = 0.143). treating each hit as a node in a graph and connecting pairs of hits separated by exactly 4 bp (the expected repeat-gap size). Connected components are found by depth-first search. The box C/D pair whose largest cluster contains the most intervals is selected, subject to a minimum of four intervals.

Each seed cluster is then extended bidirectionally using a six-level defect hierarchy applied greedily at each candidate position. The levels, tried in order, are: (1) perfect motifs at the exact expected position; (2) perfect motifs with a ±1 bp positional shift; (3) up to one mismatch in box C or box D at the exact position; (4) up to one mismatch with a ±1 bp shift; (5) perfect motifs with a spacer length of 8, 9, or 10 nt; (6) up to one mismatch with a spacer length of 8–10 nt. A candidate interval is accepted at the first level that yields a valid match, provided it falls 4–5 bp from the current cluster boundary. Extension halts when no further intervals can be added in either direction. Overlapping clusters are resolved by retaining the larger cluster.

After extension, the algorithm determines the reading frame, i.e. which gaps between consecutive intervals are loop repeats (internal to a tigRNA unit) and which are edge repeats (between consecutive units). The gaps alternate between two types, and the correct phase is inferred from base complementarity: for each of the two possible framings, the proportion of 4-nt gaps whose terminal bases are Watson-Crick or G-T wobble complements is calculated, and the framing with higher complementarity is assigned as the edge phase. Ties are broken by average pairwise sequence identity among the gap sequences. This heuristic exploits the structural observation that edge repeats form short intramolecular stems in processed tigRNAs^1^. Intervals are then paired across loop gaps to define tigRNA units. Clusters with fewer than four intervals after extension are discarded. Arrays with three or fewer assembled units are retained only if all box C and box D motifs match the reference pair perfectly; larger arrays are retained regardless.

*Stem-loop array detection.* Stem-loop arrays use a different set of box C (CCA, CGA, CTA, CAA) and box D (TG, CG) motifs, yielding eight combinations. For each combination, all perfect 14-nt intervals are identified using overlapping pattern matching. Pairs of non-overlapping hits separated by at most 15 bp are formed as candidate stem-loop units. For each pair, the stem length is measured by extending outward from both flanks one base at a time, comparing the upstream sequence to the reverse of the downstream sequence using a wobble-aware Levenshtein distance in which complementary substitutions (A-T, C-G, G-T) cost zero.

Extension continues as long as the cumulative edit distance does not exceed a leniency of 1. Only pairs with a stem length of at least 5 bp are retained. Overlapping pairs are resolved greedily in favor of the longest stem (ties broken by shortest internal gap distance). The box C/D combination producing the most valid pairs is selected, with average stem length as a tiebreaker.

The initial pair set is then expanded through four variant-search phases with progressively relaxed motif constraints and correspondingly elevated stem thresholds. Phase 1 (single-nucleotide variants of box C, correct box D, spacer = 9 nt) and phase 2 (correct box C, single-nucleotide variants of box D, spacer = 9 nt) each require a minimum stem of 8 bp. Phase 3 (variants of both box C and box D, spacer = 9 nt) and phase 4 (correct motifs, spacer = 8 or 10 nt) each require a minimum stem of 13 bp. The escalating stem thresholds compensate for the reduced specificity of relaxed motif matching, ensuring that only structurally well-supported

### Phylogenetic tree construction

Multiple sequence alignment was generated following our previously described protocol^1^. Briefly, NOP domain of TaTasR ^1^ was structurally aligned to each Tas representative (70% of sequence identity) using DALI^22,23^. All pairwise alignments were combined into a multiple sequence alignment using NOP TaTasR ungapped positions. A maximum-likelihood phylogeny was inferred from this masked alignment using IQ-TREE 3.0.1 with automatic model selection (ModelFinder; best-fit model BLOSUM62+R10). Branch support was assessed with 1,000 ultrafast bootstrap and 1,000 SH-aLRT replicates.

Tree was visualized in iTOL^24,25^ (Fig. 2a). To assess catalytic dyad conservation, Tas proteins were structurally aligned to SpTasH NOP domain (PDB:9W03) ^4^. Using conserved dyad positions at Y289 and H296, we identified catalytic motifs within dual repeat systems (Extended Data Fig. 1a).

### Structural insertion determination

To characterize structural variability, we used the TaTasR NOP domain as a reference scaffold for measuring structural extensions across the Tas70 group. Following DALI structural alignment, we systematically partitioned the NOP core into defined segments comprising the coiled-coil helices (K109-D138; A142-H168), tip (R139-D141), wedge (G210-K230), and C-terminal domain (P169-N322; insertion1: Q247-T249; insertion2: V267-S269) (Fig. 2b). We then quantified insertions within these regions by counting residues occupying gapped positions relative to the TaTasR seed. This provided a systematic measurement of diversity within the tip, wedge, and C-terminal Insertions 1 and 2 (Fig. 2a-c, Extended Data Fig. 2b).

For each Tas protein of the tree, up to 10 kb of flanking genomic context was extracted both side of the Tas coding sequence where available; because many source contigs are shorter than this window, the recovered sequence had a median length of 5,374 bp (39.9 Mb across 4,817 loci). Source annotations were retrieved for all accessions from NCBI Datasets, ENA and MGnify. Organism-level taxonomy was recoverable only for the non-metagenomic subset (366/4,817, 7.6%), as metagenomic accessions identify a sample rather than an organism; these assignments took precedence over prediction. Host taxonomy for the remainder was predicted with MMseqs2 against GTDB (R232) using easy-taxonomy --search-type 2 --tax-lineage 1, which assigns a per-contig last-common-ancestor over predicted ORFs ^26–28^. Viral and proviral origin was predicted with geNomad v1.12.0 (database v1.9) ^29^ using end-to-end -- enable-score-calibration. Because geNomad’s false-discovery filter excludes sequences carrying substantial viral signal (783 loci without a call have calibrated virus scores ≥ 0.5), loci were partitioned on the calibrated score rather than the filtered calls: viral (a geNomad call, or score ≥ 0.5), ambiguous (0.2–0.5), cellular (< 0.2). CPR was defined as GTDB phylum *Patescibacteriota* ^30,31^ unioned with accessions carrying an NCBI Patescibacteria-group annotation (n = 431). Sub-classification used the same lineage at class and order level, resolved for 93% and 87% of CPR loci respectively; family and genus were too fragmented to use (Supplementary Table S1).

## Data Availability

All TIGR annotations can be explored interactively via the web browser which will be made publicly available upon publication. Full CSV annotation files for dual-repeat and stem-loop annotations will be made available on zenodo upon publication.

The atomic coordinates and cryo-EM density maps for the PateTasR and CsTasR complexes have been deposited in the Protein Data Bank (PDB) and the Electron Microscopy Data Bank (EMDB) and will be available upon publication.

## Code Availability

TIGRFinder is freely available as open-source software and will be available on github upon publication. An archived version of the code is deposited on Zenodo and will be available upon publication.

## Acknowledgements

We thank all members of the Zhang lab for support and helpful discussions. We thank H. Kuang, B. Wang, and W. J. Rice at NYU Langone Health’s Cryo-Electron Microscopy Laboratory (RRID: SCR_019202) for help with microscope operation and data collection; S. Sterling and J. Podgorski for the maintenance and operation of the MIT.nano cryo-EM facility, established in part with financial support from the Arnold and Mabel Beckman Foundation. F.Z. was supported by the Howard Hughes Medical Institute; Yang Tan Collective at MIT, the K. Lisa Yang and Hock E. Tan Molecular Therapeutics Center; Broad Institute Programmable Therapeutics Donors; and the BT Charitable Foundation.

## Author Contributions

P.X., L.Y.L., G.F. and F.Z. designed the study. P.X., S. Z-T., N. Q-O., and L.E. conducted structural and molecular analyses. L.Y.L., G.F., A.Z., S.K., and D. F-S. performed computational analyses. R.M. and F.Z. acquired funding. G.F. supervised the computational work. R.M. and F.Z. supervised the work. P.X., L.Y.L., G.F., and R.M. wrote the original draft; G.F., R.M. and F.Z. revised, reviewed and edited the draft.

## Extended Data

### Extended Data Figure Legends

**Extended Data Figure 1.**
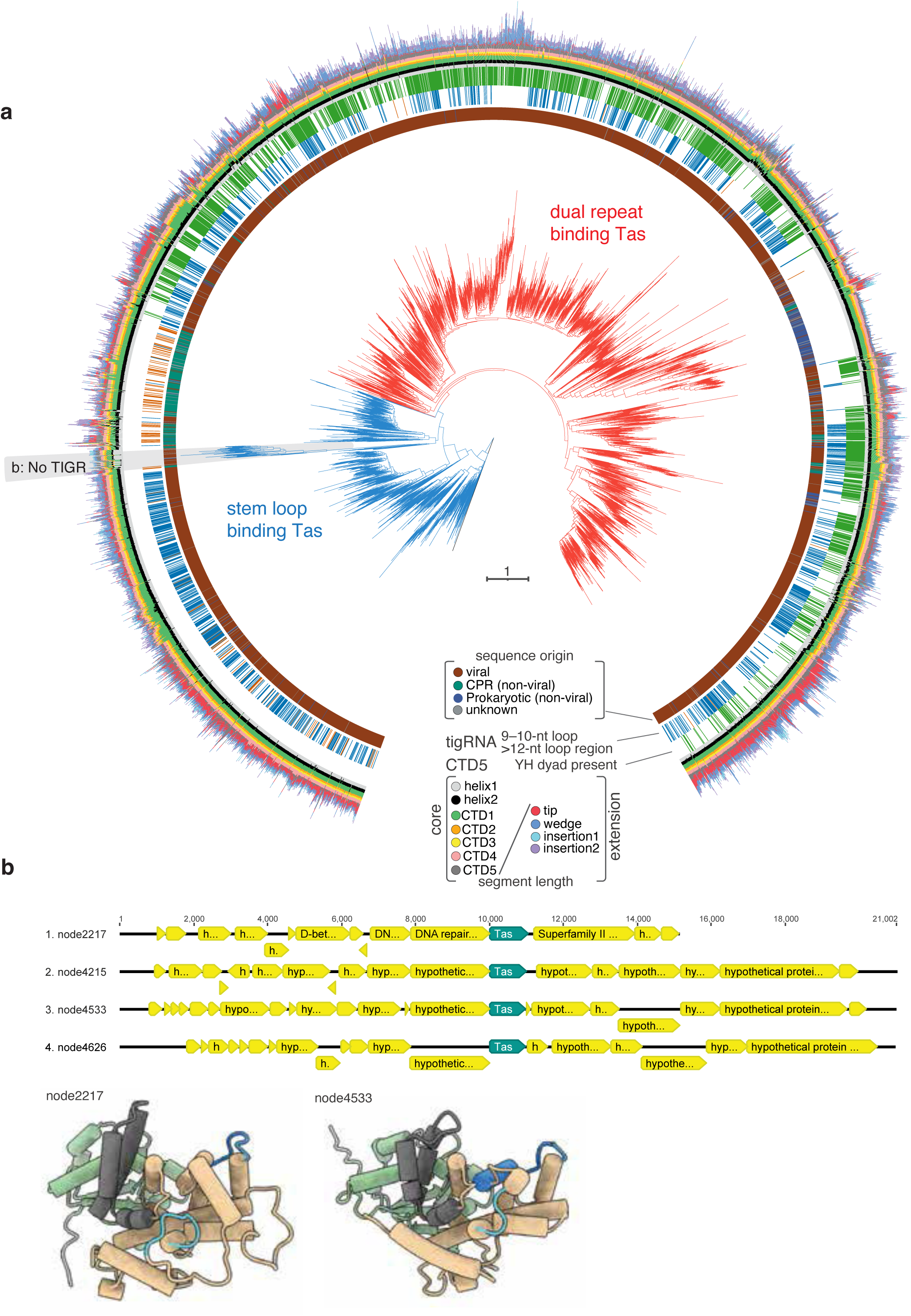
Phylogenetic and genomic characterization of TIGR systems reveals structural and architectural diversity across dual-repeat and divergent clades. **(a)** Expanded phylogenetic tree of all Tas protein representatives (clustered at 70% of sequence identity). Inner to outer rings show: tigRNA loop region length (blue for typical 9-10 nt, brown for extended >12 nt), presence of Y/H dyad, and NOP domain extensions colored by their internal location (same as Fig. 2a). Divergent *tas* clade with no coiled-coil dimerization domain and no predicted array is highlighted. **(b)** Example loci of divergent *tas* clade with node IDs. *Tas* coding sequence in turquoise, other predicted coding sequences in yellow. Representative structures as predicted by AlphaFold are shown, colored the same as in Fig. 2b.

**Extended Data Figure 2.**
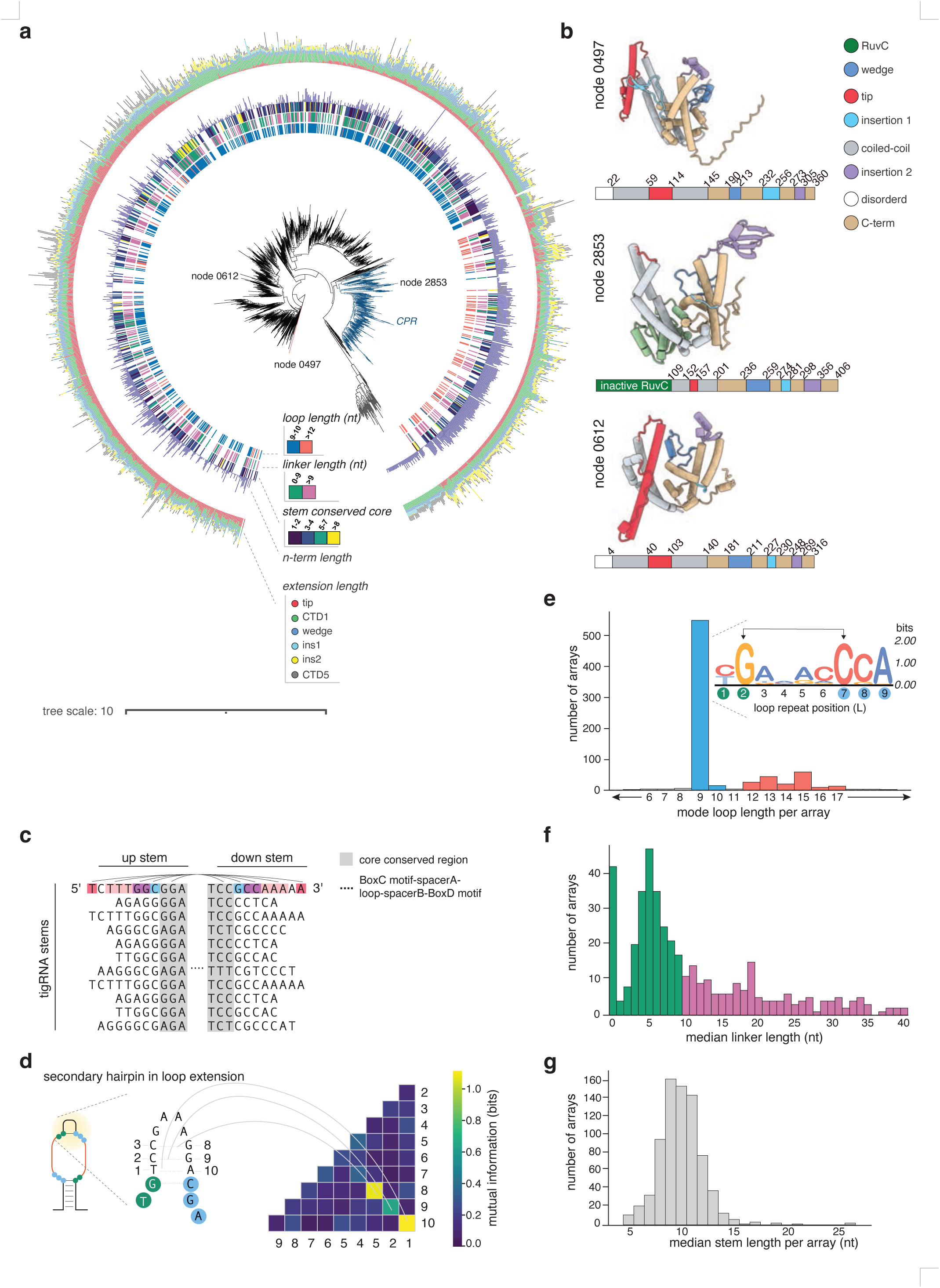
Structural and sequence diversity of tigRNA components and their covariation with Tas protein architecture. **(a)** Phylogenetic stem-loop tree (same in Fig. 2a) depicting tigRNA component diversity coevolution with tas extension lengths. CPR clade branches are labeled in dark-blue. Tas proteins with unique secondary structures in domains are labeled, see (b). Tree annotation rings from innermost to outermost: mode tigRNA loop length (9-10 nt in blue, >12 nt in orange), median linker length (0-9 nt in green, >9 in purple), the number of conserved stem nucleotides (1-2 nt in purple, 3-4 nt in dark-blue, 5-7 nt in dark green, >8 nt in yellow). Specifically, we define a “diversity ratio” to be the number of unique sequences divided by the total number of units in an array. Then, for each stem-loop array, the stem conserved core length is the longest sequence of nucleotides from the C (D) motif for the upstream (downstream) stem to reach a diversity ratio greater than 0.5. Other Nop domain extension annotations are the same as in Fig. 2a. **(b)** Selected Tas with defined secondary structures in tip, insertion 1, and/or insertion 2 components. Coloring same in Fig. 2d. **(c)** Example of a stem conserved core. The upstream and downstream stems are shown, with the main functional tigRNA unit in “…”. Conserved stem core is highlighted in gray. Complementarity shown with matching colors. **(d)** Example of extended loop region. Mutual information table confirms covariation (conserved complementarity) for extended loops to extend as a secondary stem. D motif shown in purple, C motif shown in cyan. **(e)** Distribution of mode loop length per stem-loop array. There is a peak at 9nt loop lengths, with a sequence motif of all representative 9nt stem loop lengths shown, with height representing information content in bits. Conserved structural complementarity is shown with thin arrows. D motif positions highlighted in green, C motif positions highlighted in cyan. Blue and orange correspond to colors of loop length ring on tree in (a). **(f)** Distribution of median linker length per stem-loop array. Stem-loop arrays with overlapping stems are defined to have 0 linker length. Green and purple correspond to the colors of the linker length ring on the tree in (a). **(g)** Distribution of median stem length per stem-loop array.

**Extended Data Figure 3.**
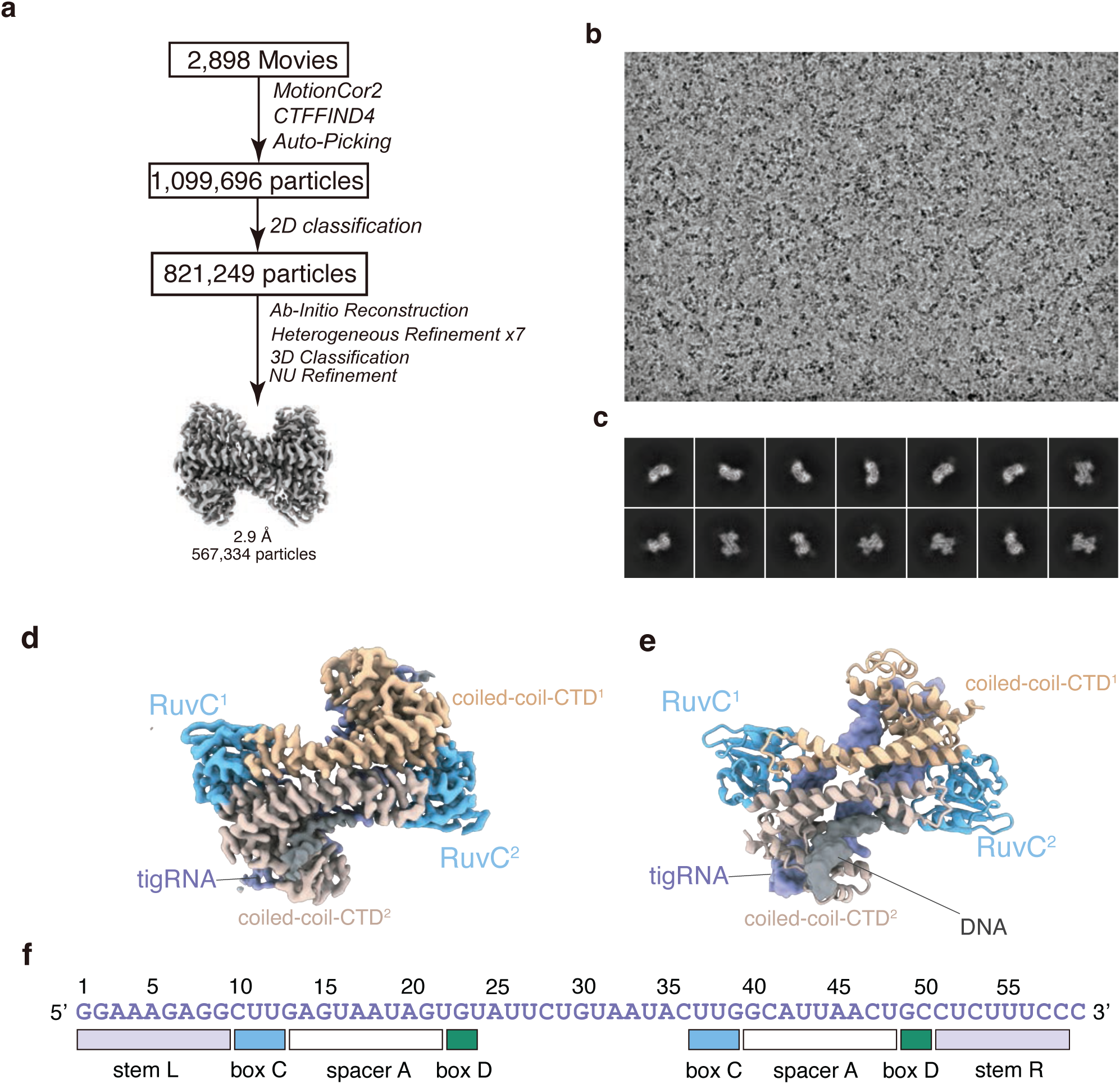
Cryo-EM data processing of stem-loop PateTasR. **(a)** Flow chart of cryo-EM data analysis of *Patescibacteria group bacterium* TasR (PateTasR). **(b)** Representative cryo-EM image of PateTasR from a dataset of 2,898 movies. **(c)** Representative 2D averages of PateTasR. **(d)** Cryo-EM density map of the PateTasR complex. Color coding: RuvC domain, blue; coiled-coil scaffold, gray; C-terminal domain (CTD), tan; tigRNA, purple; target substrate, black. **(e)** Atomic model of the PateTasR complex. **(f)** Sequence and Box C/D motifs of the PateTasR tigRNA.

**Extended Data Figure 4.**
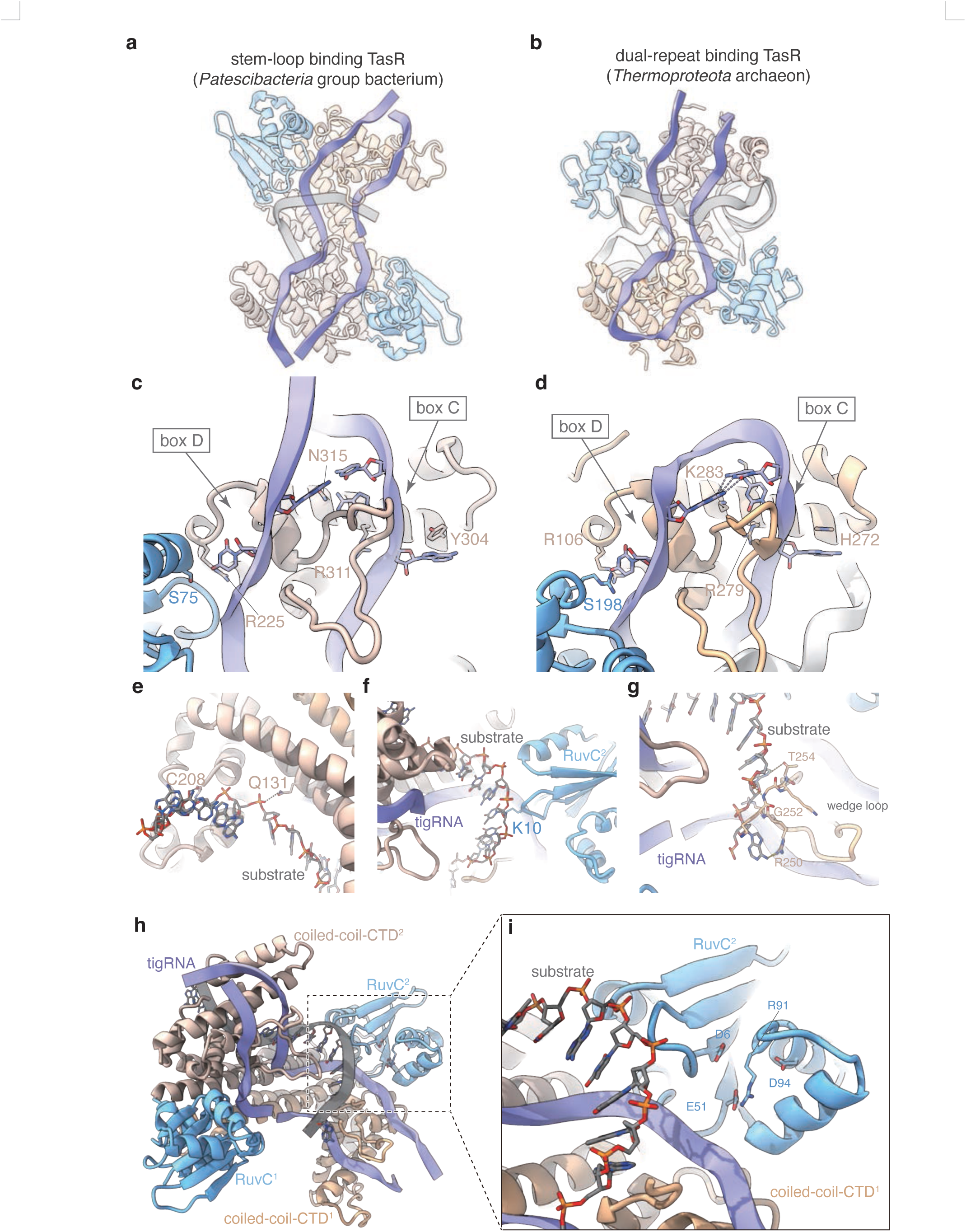
Cryo-EM structures and atomic models of stem-loop PateTasR and CsTasR complexes. **(a)** Cryo-EM density map of the *Patescibacteria* group bacterium TasR (PateTasR) complex. Color coding: RuvC domain, blue; coiled-coil scaffold, gray; C-terminal domain (CTD), tan; tigRNA, purple; target substrate, black. **(b)** Atomic model of the PateTasR complex. **(c)** The tigRNA sequence and Box C/D motif annotation of PateTasR. **(d)** Cryo-EM density map of the *Candidatus Staskawiczbacteria bacterium* TasR (CsTasR)- tigRNA-target DNA complex. Color coding: RuvC domain, blue; coiled-coil scaffold, gray; CTD, tan; tigRNA, purple. **(e)** Atomic model of the CsTasR-tigRNA-target DNA complex. **(f)** Secondary (2D) structure of the CsTasR tigRNA. **(g)** AlphaFold 3 model of BlTasA; the wedge loop is shown overlapping with two tigRNA coding regions. **(h)** Structure of the CsTasR RuvC domain; two catalytic site residues are substituted with arginine (R) and valine (V).

**Extended Data Figure 5.**
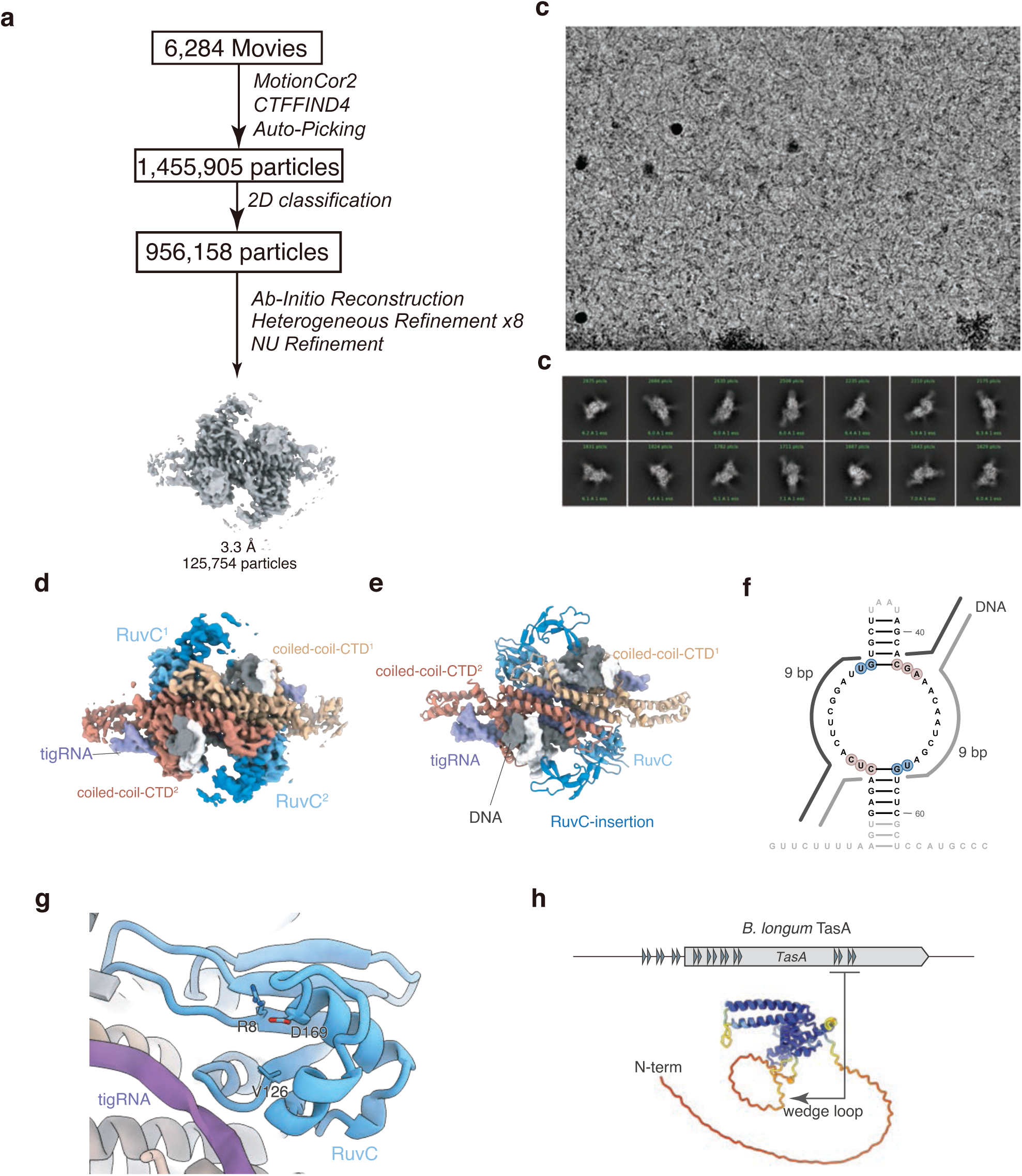
Interactions of PateTasR with tigRNA and structural comparison with dual-repeat TaTasR. **(a)** Overall structure of PateTasR in top view; the tigRNA exhibits a figure-eight conformation with stem structures at both ends. Color coding: RuvC domain, blue; coiled-coil scaffold, gray; C-terminal domain (CTD), tan; tigRNA, purple; target substrate, black. **(b)** Overall structure of dual-repeat TaTasR in top view; the tigRNA features a similar figure-eight architecture. **(c, d)** Structural comparison of the Box C/D motifs between stem-loop PateTasR (c) and dual-repeat TaTasR (d). **(e–g)** Detailed views of interactions between the substrate and specific protein regions: the coiled-coil (CC) domain (e), the RuvC domain (f), and the wedge loop (g). **(h, i)** Detailed view of the PateTasR RuvC catalytic site. The catalytic residues are physically obstructed by an arginine (R) residue.

**Extended Data Figure 6.**
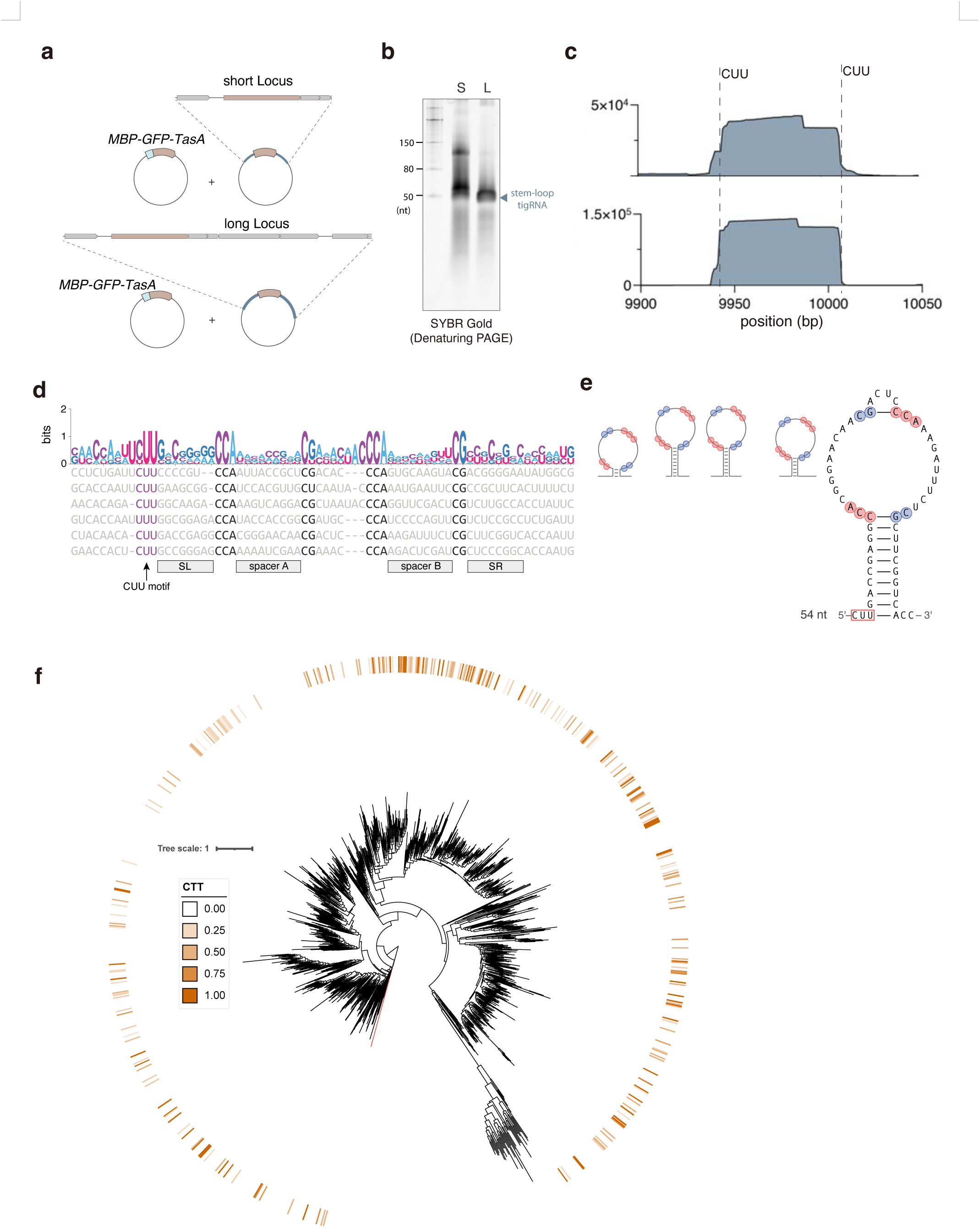
Small RNA-seq reveals that extended genomic context facilitates tigRNA maturation and identifies a conserved CUU motif as a potential cleavage site. **(a)** Schematic of BlTasA co-expression strategies using either a compact or an extended genomic locus; the extended locus includes several coding sequences (CDS) downstream of *tasA*. **(b)** Denaturing PAGE analysis of purified RNPs from the compact and extended genomic loci. **(c)** RNA-seq tracks displaying the CUU motif at the boundaries of tigRNAs. **(d)** Sequence alignment of BlTasA tigRNAs; the conserved CUU motif is annotated across sequences. **(e)** Secondary (2D) structure of tigRNA; the CUU motif is located at the base of the stem. **(f)** Stem-loop tree (same as Fig. 2a) with annotations showing proportion of extended linkers (including last two nucleotides of preceding downstream stem and first two nucleotides of following upstream stem) that are less than 20nt that contain a CUU motif.

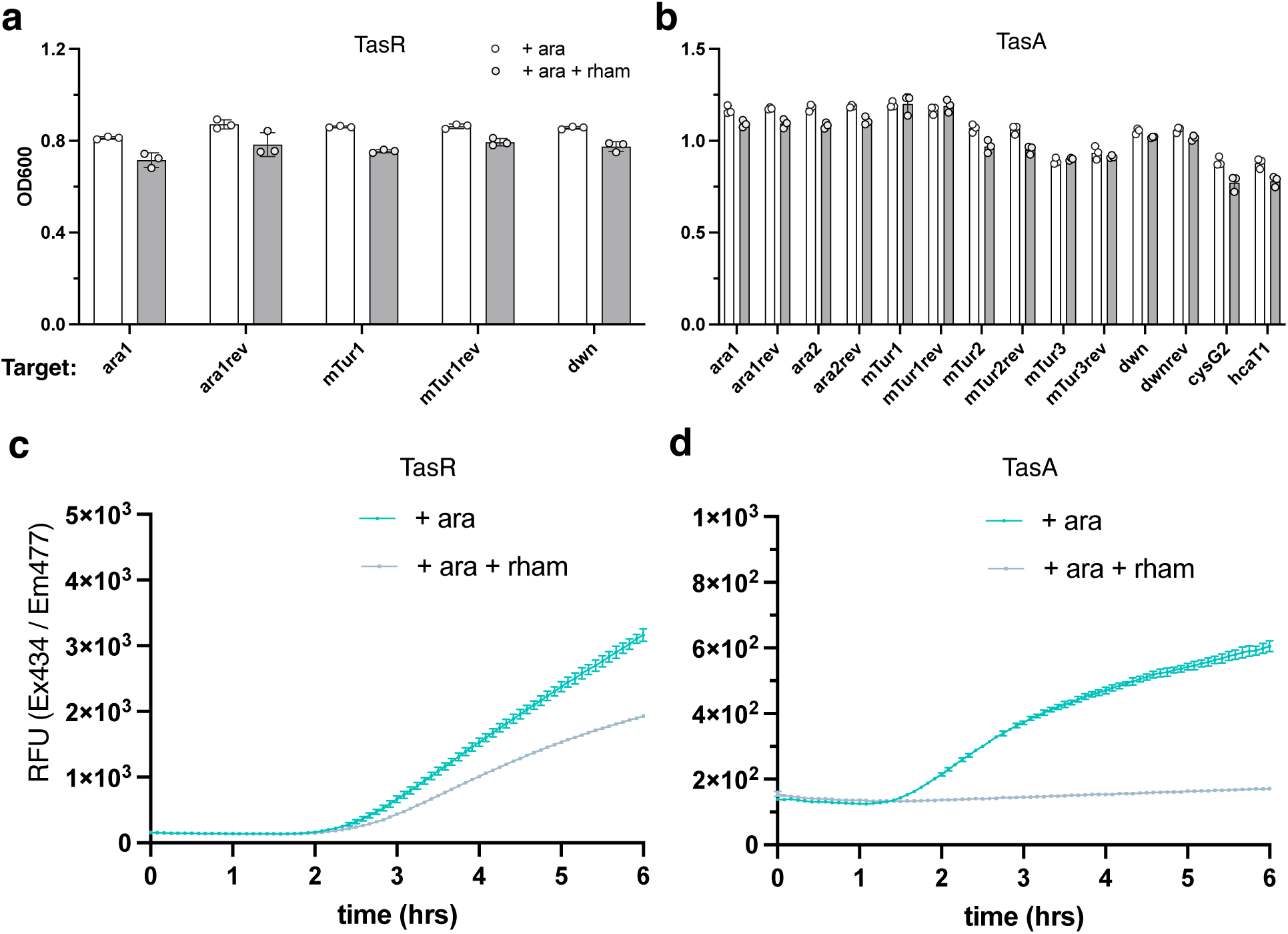

**Supplementary Table**. All experimental loci and associated sequences used in this study. Node ID (from phylogenetic tree) to full contig ID mapping included. File will be available upon publication.

